# AuPairWise: a method to estimate RNA-seq replicability through co-expression

**DOI:** 10.1101/044669

**Authors:** Sara Ballouz, Jesse Gillis

## Abstract

In addition to detecting novel transcripts and higher dynamic range, a principal claim for RNA-sequencing has been greater replicability, typically measured in sample-sample correlations of gene expression levels. Through a re-analysis of ENCODE data, we show that replicability of transcript abundances will provide misleading estimates of the replicability of conditional variation in transcript abundances (i.e., most expression experiments). Heuristics which implicitly address this problem have emerged in quality control measures to obtain ‘good’ differential expression results. However, these methods involve strict filters such as discarding low expressing genes or using technical replicates to remove discordant transcripts, and are costly or simply ad hoc. As an alternative, we model gene-level replicability of differential activity using co-expressing genes. We find that sets of housekeeping interactions provide a sensitive means of estimating the replicability of expression changes, where the co-expressing pair can be regarded as pseudo-replicates of one another. We model the effects of noise that perturbs a gene’s expression within its usual distribution of values and show that perturbing expression by only 5% within that range is readily detectable (AUROC~0.73). We have made our method available as a set of easily implemented R scripts.

**Author Summary:** RNA-sequencing has become a popular means to detect the expression levels of genes. However, quality control is still challenging, requiring both extreme measures and rules which are set in stone from extensive previous analysis. Instead of relying on these rules, we show that co-expression can be used to measure biological replicability with extremely high precision. Co-expression is a well-studied phenomenon, in which two genes that are known to form a functional unit are also expressed at similar levels, and change in similar ways across conditions. Using this concept, we can detect how well an experiment replicates by measuring how well it has retained the co-expression pattern across defined gene-pairs. We do this by measuring how easy it is to detect a sample to which some noise has been added. We show this method is a useful tool for quality control.

## Background

Recent analyses of RNA-seq have emphasized the value added by experimental designs with more samples, permitting better estimation of biological and technical variability [1-4]. Unfortunately, this does not automatically translate into an experimental design decision for more replicates (biological or technical), largely due to the relatively high cost of RNA-seq, which can range anywhere from 10c to $10 per Mbp [5]. Study design is further complicated by complex dependencies on the platform chosen, library preparation and normalization methods; all these factors affect the concordance of downstream differential expression analysis, as a recent set of large scale studies have systematically demonstrated [5-9]. While researchers have to be aware of the pitfalls that can arise when designing their experiment, they receive only modest feedback in the experimental data itself as to whether their choices were successful. While many optimized protocols or best practices exist, there are few means for assessing the quality of a given transcriptomic study, particularly outside of purely technical concerns and where focusing on novel biology.

Systematic and large scale studies assess new technologies for appropriate guidelines and best practices at great cost, time and effort. When microarrays began to be used for drug development, the Food and Drug Agency (FDA) set up the MicroArray Quality Control (MAQC) for general standards of practice in order to assure high quality and replicability [10]. In a similar fashion for RNA-seq, Sequencing Quality Control (SEQC or MAQC-III) and the Association of Biomolecular Resource Facilities (ABRF) were tasked to determine a set of gold-standard pipelines, protocols and metrics for assessment [5,9].

Extensive work and funding were put into these efforts: evaluating terabytes of data, across numerous preparation conditions, platforms, sample types, experiment sizes, read depths, computational quantification methods, statistical measurements and other diagnostics. The projects successfully identified the strengths and weaknesses of various aspects of RNA-seq but, alarmingly, could not generalize these findings to a consensus gold-standard method or technique. Different platforms gave different biases, depending on the chemistry of the reactions, length of the reads, the amplification of the sample and method of degradation of ribosomal RNA, to name a few. These are reflected in underlying sequence-based biases from GC content, genome size and gene size.

Many of these properties are addressed directly in quality control practices which have a strong history in microarray analyses and can be applied to RNA-seq without too much loss of generality. A notable example is batch detection and correction, long identified as critical in microarrays (e.g., [11]), and for which methods have been adapted for use in RNA-seq [7,12]. Technical quality control metrics attached to the samples, such as RNA integrity [13], read coverage, GC content, etc. (e.g, RSeQC [14], RNA-SeQC [15]), may also be applied without alteration to RNA-seq analyses, even if the effect of such biases may be technologically dependent. Novel quality control assessments for RNA-seq have also been developed or adapted such as the irreproducible discovery rate (IDR), initially developed for CHIP-seq data [16], and then applied by ENCODE to RNA-seq data [17]. Competing statistical approaches to detect replicable signals (e.g., SERE [18]) are also in use. In all these cases, the quality control is more focused on identifying or correcting purely statistical properties without exploiting biology in any way. The latter task is challenging precisely because the biology depends on the exact experiment being performed.

Similarly, spike-ins of known concentrations of RNA, such as those of the External RNA Control Consortium (ERCC) [19], can be used for quality control in specific cases and to identify technical artifacts but they are useful precisely because they are independent of the biology of the experiment. Even so, their results suggest variation in expression from experiment to experiment that is large enough that there is no way to universally ‘normalize away’ our biases in a one-size-fits-all manner [7]. Instead of standardizing reads, filters are recommended to remove unwanted variation, which in the case of the recommendations from SEQC/MAQC-III, includes discarding a third of the genes with low expression levels and those outside a fold change (FC) threshold between samples (which was algorithm dependent, log_2_ FC ~1-2). But these are general rules whose relevance must vary from case to case. How well do these heuristics generalize to carefully designed experiments conducted in an entirely different biological system?

In most individual labs, there will be little to no data to assess this question directly. Few labs can afford experimental designs which use samples purely as replicates for quality control. This is unfortunate, since often the experimental design is specifically intended to address noise in the biological system. There is no easy and systematic way of seeing whether this worked, outside of the experiment giving expected results, but it is unexpected results that are particularly valuable. Ideally, the results that are used for validation can provide a finely calibrated sense of performance without impinging on the capacity for novel discovery, but this will be challenging since the same data will likely be used in both cases. As a consequence of this, assessment of RNA-seq efficacy in any targeted experiment is typically difficult, relying on experts in the particular experimental system finding the results plausible (but not too obvious). The few truly generally applicable quality controls are extremely coarse, and focus more on technical properties, such as read depth, coverage, and related measures.

Ideally, each lab could conduct their own RNA-seq quality control, relying either on costly replicates, or some sub-class of ‘expected’ result which does not diminish the freedom to obtain other unexpected results. Our approach to this problem is to focus on modelling the behavior of tightly co-expressed genes. Gene-pairs which are very tightly co-expressed can be thought of as a form of a pseudo-replicate and have the advantage of reflecting a broadly applicable biological property orthogonal to the tested variation in most studies, which are usually based on differential expression. Of course, to the extent we rely on ‘knowing’ pairs of genes are co-expressed to determine experimental success, we diminish our ability to find we were mistaken about this property. To deal with this issue, our analysis relies on weak relationships across many genes but no strong knowledge of any single gene and only a tiny fraction of gene-gene-pairs or ‘housekeeping interactions’[20].

In this paper, we demonstrate a straightforward but surprisingly powerful method, which we have named AuPairWise, to measure replicability by modelling the effects of noise within observed co-expression. Because of the breadth and extent of co-expression, it provides a sensitive, yet general means of performing a quality control. We argue this in roughly three steps. We begin our analysis by performing some sample and replicate-based analyses of RNA-seq quality to establish general properties. Second, we show how common heuristics for quality control appear in a co-expression based framework. Finally we provide a means of directly quantifying this by disruption of known co-expression pairs using a model based on empirical distributions of expression data.

By providing a direct quality control measure, we hope to make it straightforward to customize experiments to do better than a general heuristic - such as discarding all low-expressing genes – would normally permit. We make our software available in a convenient form for bioinformatic use (https://github.com/sarbal/AuPairWise).

## Results

### PART I: Reviewing replicability

#### Two measures of expression replicability in RNA-seq

Before assessing replicability, it’s necessary to provide a sense of the meaning which we attach to the term. Broadly, replicability in RNA-seq is used in two quite different ways, one focusing on the replicability of estimates of mean gene expression (sample-level replicability) and the other on variation in gene expression (gene-level replicability). The earliest method is to assess whether a given pair of samples is well correlated across its transcripts or genes. This measures whether genes have similar expression in the two samples and by this standard, all RNA-seq has high enough replicability that we never need worry. Unfortunately, this doesn’t map to a very experimentally useful measure since most experiments are concerned with a gene’s variation in expression under changing conditions (i.e., for differential expression). To see why these two forms of replicability do not align, consider an early reference experiment [21] which reported high replicability in a given pair of samples from the liver, as measured by a correlation across transcripts of Spearman’s *r_s_*=0.976 (p<1e-15, Fisher’s transformation) (Fig. 1A). But this correlation is partially a property of transcript abundances having an approximate transcript-specific range defined regardless of tissue. As we can see, the correlation comparing the liver and the kidney is also very high (Fig. 1B), and indeed, variation from one experiment [22] to another is sufficient to make correlations across tissues within an experiment appear much greater than those within tissues but across experiments (Fig. 1C).

**Fig. 1.**
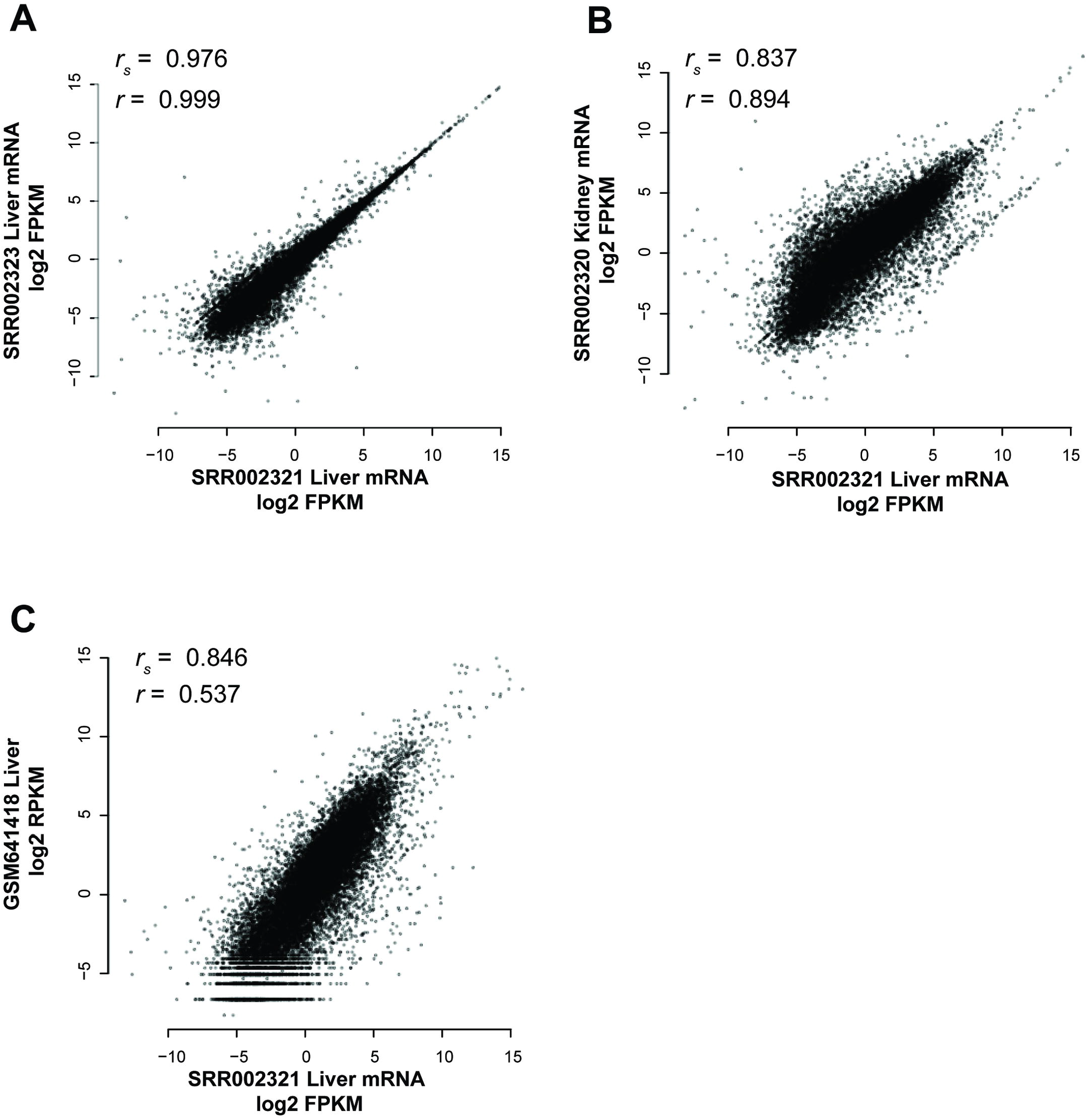
Individual genes have similar expression levels in many tissues. Samples replicate one another to some degree, regardless of the conditions under which they are measured, i.e., whether it is actually a biological replicate or not.(A) Liver expression levels within the same experiment are very highly correlated (Spearman’s *r_s_* =0.976, Pearson’s *r*=0.999). (B) Liver expression is moderately correlated to kidney expression within the same experiment (Spearman’s *r_s_* =0.837, Pearson’s *r*=0.894). (C) Liver expression levels in two different experiments are less well correlated than that between tissues (Spearman’s *r_s_* =0.846, Pearson’s *r*=0.537).

Rather than this sense of replicability, we are primarily concerned with the degree to which changes in expression for a given gene are replicable (i.e., gene-level replicability or self-correlation). While the importance of replicability in terms of differential expression has been appreciated from the first MAQC forward [23], it is difficult to generalize from differences due to specific conditions (e.g., tissues vs disease) to make cross-experiment comparisons characterizing quality control. It is this differential expression sense of replicability that is useful in most biological experiments, but the problem in even assessing it within a given experiment is that it requires enough samples to at least partially replicate the entire experiment. Additionally, the degree to which genes show replicable differences will depend in part on the degree to which they show condition-dependent variability at all.

#### A contradiction between the replicability measures

As a brief experiment to further motivate a different metric of replicability, consider data for which we have a replicate for each expression measurement across multiple conditions for each gene; essentially, extending Fig. 1A to even more conditions and plotting all the points at once. We show this using reference data from ENCODE in Fig. 2, where each point represents a gene under a given condition with x-values being the initial expression measure and y-values being the replicate measurement (for that gene under that condition, see Materials and Methods section “Measuring replicability: correlations and sample estimates). First, we show the gene-gene correlations for each sample (Fig. 2A, each sample is a color, we show 9 of the 18 samples for clarity) and then the overlay of these as a replicate-replicate plot (Fig. 2B). Clearly, there is a high level of correlation between the measurements of expression. We can display specific genes in that same plot by highlighting them; each with quite distinct dynamic ranges (colored in Fig. 2C and D). Because genes will only vary within their own dynamic range, collectively measuring replicability ignores the gene specific variation. A given gene’s measured expression could be well correlated with the same gene when measured in the replicate data (the subset highlighted in Fig. 2C) or poorly correlated (subset highlighted in Fig. 2D), but the correlation across genes for a given condition would not reveal it: the fact gene A is higher in expression than gene B is not at all condition dependent (because of the disjoint dynamic ranges). Thus, the more genes have an approximate set point in expression, the more trivially replicable the correlation of transcript abundances between two samples regardless of condition, and the less the two measures of replicability have any relation. Ironically, one of the chief claims made of RNA-seq – that it has high dynamic range – would only increase this tendency, since it provides more ‘space’ for each transcript to occupy a separate expression domain, yielding perfect but meaningless sample-sample correlations. Note that here, as in the rest of this paper, we are concerned solely with gene-level mapping and use ‘transcript abundance’ to refer simply to a given gene’s expression. Transcript variability for a given gene is, of course, a topic of great interest, but are challenging in their own right. We focus here on establishing more basic properties first.

**Fig. 2.**
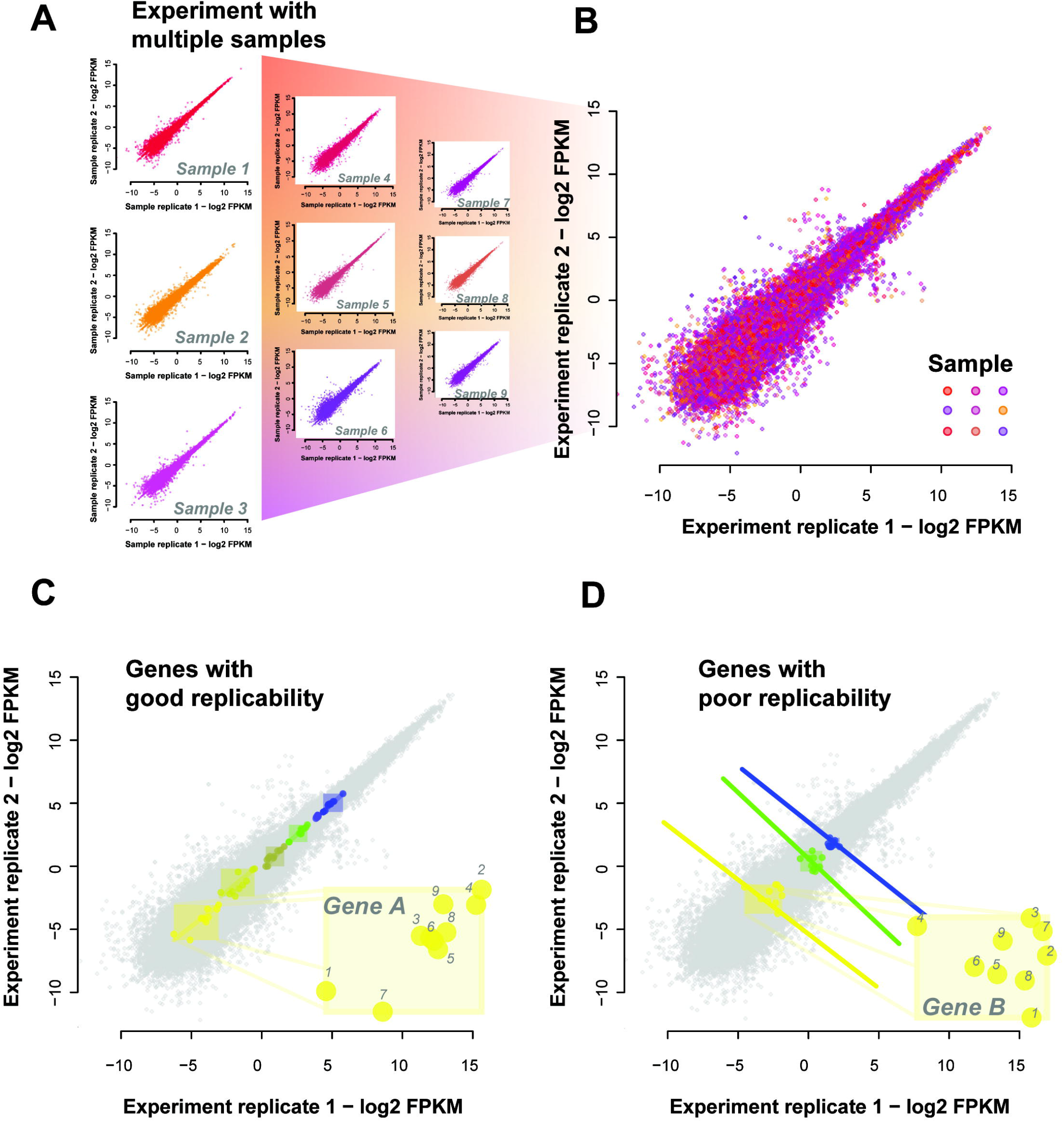
Simpson’s paradox in expression data. (A) Sample-sample replicability plots are possible for each of the 18 conditions tested. Shown are a representative set of 9 sample-sample plots. Each point represents a gene with X and Y values being the two expression measures, i.e., original and replicate. Different samples are shown in different colors. (B) We overlay all these individual sample-sample plots onto the same axes to give the aggregate view of expression across samples and their replicates. This allows us to determine replication of gene variation across samples. (C) Some genes replicate well; changing their expression in a consistent way across samples and their replicates. We have highlighted genes which are positively correlated with their replicate across the samples and also have no overlapping dynamic ranges. Note that sample-sample correlations across this set of genes would be high even if the labels identifying samples were permuted (inset of “Gene A” shows labelled samples). (D) There are also genes which are negatively correlated with their replicates across samples (inset of “Gene B” shows labelled samples). And because they also do not have overlapping dynamic ranges the sample-sample correlation across these genes would remain high for any given sample pair.

### PART II: Refining replicability through expression co-variation

#### Defining ‘replicability’ when measuring expression variation

Rather than measuring dynamic range by the number of genes differentiable by their expression within some range, we really need to know the number of conditions differentiable by the expression of each gene. That is, a gene has a ‘good’ precision over its dynamic range when change in its expression provides information about the condition under which it was measured. In future, we will use ‘replicate’ to refer exclusively to gene-level replication (or self-correlation), in which we consider whether a gene’s variation from condition to condition is replicable when the same conditions are re-measured. This is similar to the idea of checking if genes are reproducibly being detected as differentially expressed, i.e., if genes are DE in both replicates, but we are measuring correlations and not fold changes.

To assess this for each gene, we would like a range of conditions which we expect that gene to vary under, i.e., different tissues, varying time points or treatments. We would also like replicates for each of those conditions. With these, we can measure the degree to which, for each gene, its variation is distinguishable. One experiment that fits this general design was ENCODE’s RNA-seq profiling experiment (GSE35584), which looked at various biological cell lines under multiple treatments. We take ENCODE’s own labelling of the data into two batches to define the grouping of the data into sets of samples. ENCODE performed a number of perturbations likely to be genome-wide in effect. In this case, if a gene’s changed expression is meaningful, it should be correlated with its replicate across the range of conditions. However, not all genes are expected to show differential activity, and particularly if they are only modestly perturbed. For each gene, we can measure whether it does, in fact, show variation that is consistent when the same condition is measured (across all conditions). In the ENCODE data (20,635 genes detected), we find after multiple test correction that over 16,000 genes were significantly correlated (q<0.05, Fisher’s transformation, Holm-Bonferroni corrected) with their replicates across the samples (Fig. 3A), giving us a coverage of approximately ~78% of the genes in this experiment (77.7% - Student’s T, or 78.2% - Fisher’s transformation).

**Fig. 3.**
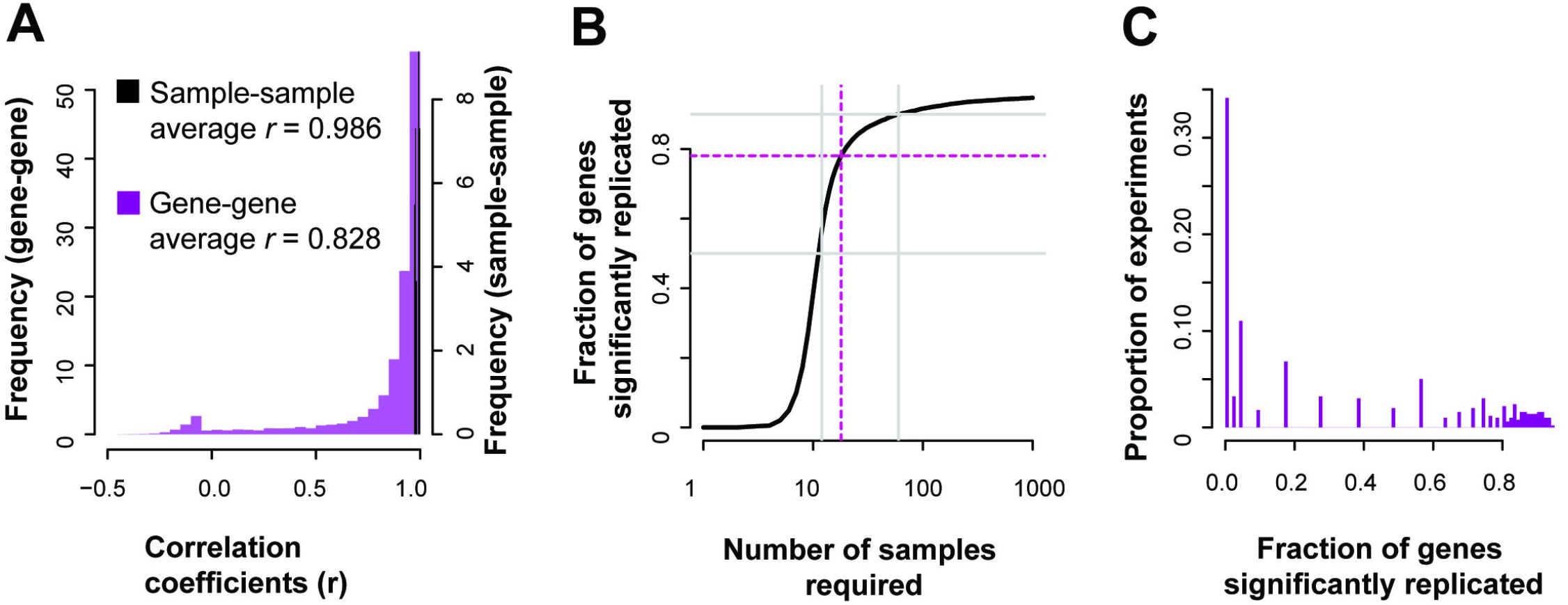
Gene expression replication across the genome. (A) In the reference ENCODE data, many genes are well correlated with their replicate across the range of conditions, but the distribution has a heavy negative tail. (B) For the same example experiment, almost 80% of the genes are significantly (q<0.05) replicated, as characterized by the correlation between gene replicates across conditions. (C) Across experiments in GEO, only a minority (~34%) are likely to be able to produce replicable results for more than half of the typically assayed transcriptome.

If we assume this correlation distribution holds constant, we can estimate that to significantly (q<0.05) cover 90% of the transcriptome we would need to increase the number of samples to over 50, and for 95% coverage, over a hundred samples (Fig. 3B, see Materials and Methods section “Measuring replicability: correlations and sample estimates for more details). Similarly, we can assess the potential coverages of prior experiments based on their sample count, and, as a proxy, assess how replicable their results are for particular genes. Evaluating 1,451 publicly available experiments, we estimate that many experiments only have enough samples to produce significantly replicable results for a small fraction of the genome, with only 34% of experiments assessed having coverage above 50% (Fig. 3C). Even though the whole transcriptome is measured in these experiments, using correlation as quantification demonstrates that we are not powered to detect variation sensitively for most of the genes with low sample sizes. Of course, this is subject to the important caveat that the experiments are, indeed, expected to produce replicable variation between control and experimental conditions only for a small fraction of genes: those actively relevant to the experimental perturbation.

One potential confound in this analysis is that increasing artifacts would broadly seem likely to increase the correlation of replicates, so long as the artifacts were maintained across replicates. Of course, this would be a problem with any sort of replication (if artifacts replicate, replication doesn’t help as much). This can be controlled for since such artifacts would generically improve the correlation between a gene and all others, rather than specifically itself in the replicate data.

We calculated the correlation of each gene with all others in the replicate data to provide such reference information. We then calculated the rank of a gene’s correlation with itself versus all other genes within the experiment. If the variation in expression that a gene shows is meaningful, we would expect it to have a higher correlation to the replicate of itself than other genes. We do observe from the distribution of self-correlation (i.e., gene-level replicability as previously defined) versus correlations to other genes (i.e., co-expression, co-regulation) that genes are much better correlated to themselves in replicate data than to other genes (Fig. 4) with almost 20% even being better correlated with their measurement in the replicate data than with any other gene in the replicate data (4,024 genes). This suggests that having closely replicating gene profiles is a relatively common occurrence in high quality data.

**Fig. 4.**
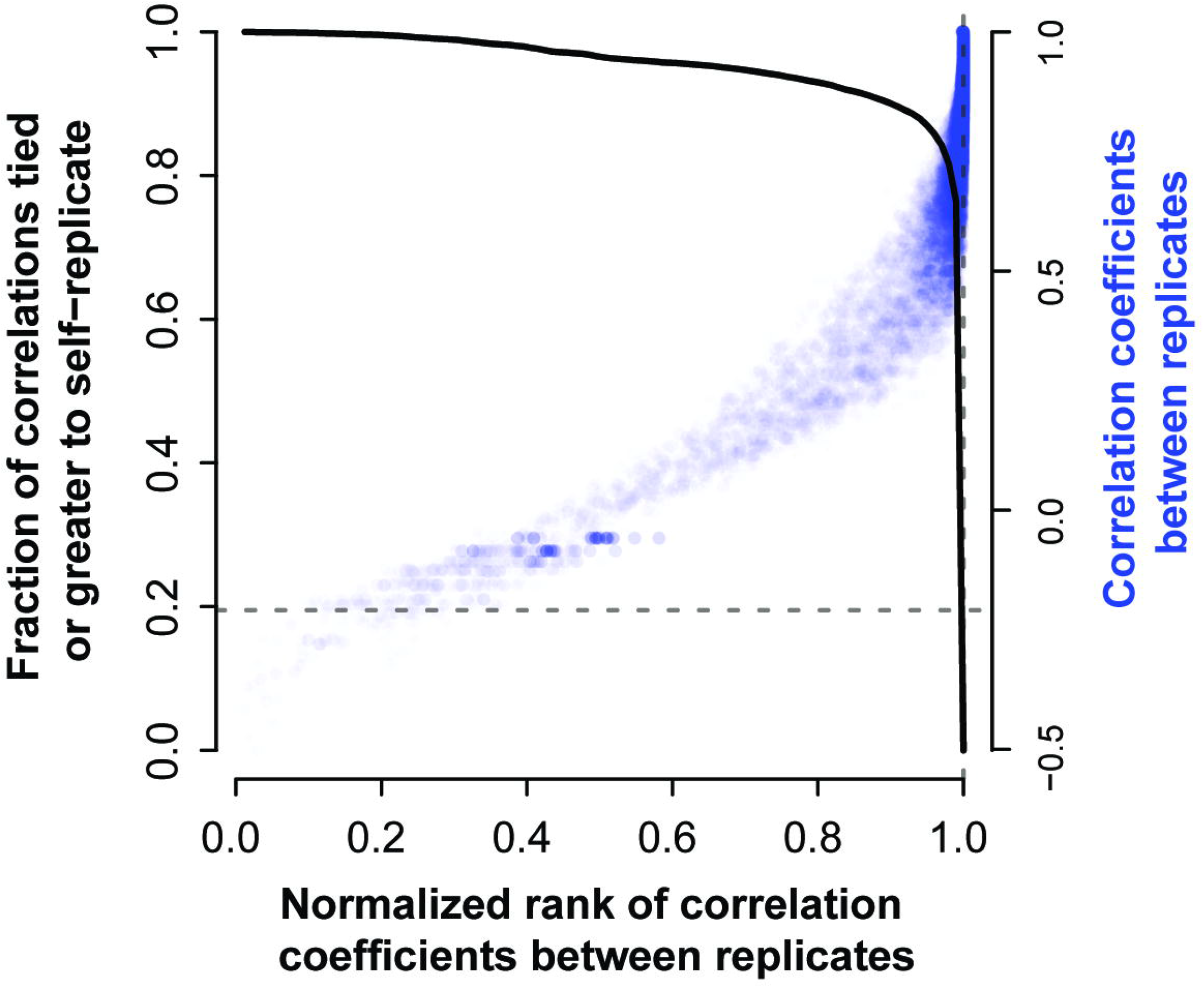
Self-correlation (replicability) and co-expression. The degree of correlation between a gene and its replicate is plotted relative to all other genes (relative rank). As would be expected, as the correlation between a gene and its replicate (across the same conditions) rises, the rank of that correlation relative to the value between the given gene and all others also rises. However, the true replicate is only most similar to the given gene in ~20% of cases (solid black line, 4,024 genes), i.e, the solid line is at 0.2 when the rank is exactly 1 (dashed lines). The steep fall off in this trend shows that most replicates are at least very highly ranked by the correct gene.

The average correlation of the genes in the experiment is high (Pearson’s r=0.97, SD 0.04). If we relax our definition slightly so that the gene has to rank its replicate within the top 10%, we see that 90% of the genes pass that threshold and that they have an average correlation of 0.9. We have used this mean value as a cutoff (Pearson’s r=0.9) to characterize ‘reasonable’ replicability for our further analysis of this study. This is more stringent than simple significant correlation; we are requiring it to be above the mean of genes with a non-parametric ‘preference’ to be correlated with themselves and close to the point nearest the top right corner (best performance). Of course, because actual biological co-expression exists (a fact we will exploit), pristine results in self-correlation are not to be expected and all downstream results are robust to any reasonable variation in this threshold. We discuss the potential role of normalization creating generic correlation across all genes in the “Robustness analysis of AuPairWise section.

#### Mapping general QC heuristics to the replicability of expression changes

We now move on to see whether this analysis of replication matches standard heuristics. The SEQC concluded that low expressing genes, equivalent to 1/3rd of the measured dynamic range of the whole experiment, and genes with lower fold changes between conditions (<log2 FC 1~2) should be discarded from analyses [9]. To assess these thresholds independently, we measure their downstream impact on replicability in our reference ENCODE data. For each gene, we plot in Fig. 5C its total mean expression (x-values) against its average fold change between conditions (y-values), and then color points according to their replicability, measured as the correlation a given gene shows with its replicates across the conditions. This allows us to visualize how the SEQC criteria align with our own definition of replicability. Overall, most of the genes with low replicability (red points in Fig. 5C) are low expressing genes and have low fold changes. Indeed, if we wish to threshold the data for particular fold changes and expression levels so that they are replicable as we have defined the term, we would end up with results similar to the SEQC guidelines (blue dashed lines in Fig. 5). Applying the fixed guidelines and removing the bottom third of expressing genes leaves us with 97% of genes showing good replicability (Pearson’s r>0.9 across replicates, p<1e-15, Fisher’s transformation [24], Fig. 5A), as compared to 61% of genes in the data overall. Likewise, poor replicability is concentrated at low fold changes in the data with irreproducible genes (Pearson’s r<0.9, Fig. 5B) having a mean fold change of 2˄0.48 and reproducible genes (Pearson’s r>0.9) having a significantly higher mean fold change of 2˄1.08 (p~2.34e-154, Wilcoxon test), similar to the threshold suggested by the SEQC experiments. One difference from the SEQC evaluation is that the irreproducible genes are not subject to their two criteria (mean expression and fold change) independently; we can see that there is some dependency between the two constraints and past a certain fold change, filtering on expression would not make a substantial difference (i.e., there are little to no genes with very high FC and low mean expression). We also note that although our definition of a “reproducible gene” here is set by a somewhat arbitrary threshold of 0.9, the mean correlation shows the same trend across the two criteria, perhaps even more distinctly (**S1 Fig**). Thus, we suggest self-correlation across conditions is a more useful and practical way of defining replicability.

**Fig. 5.**
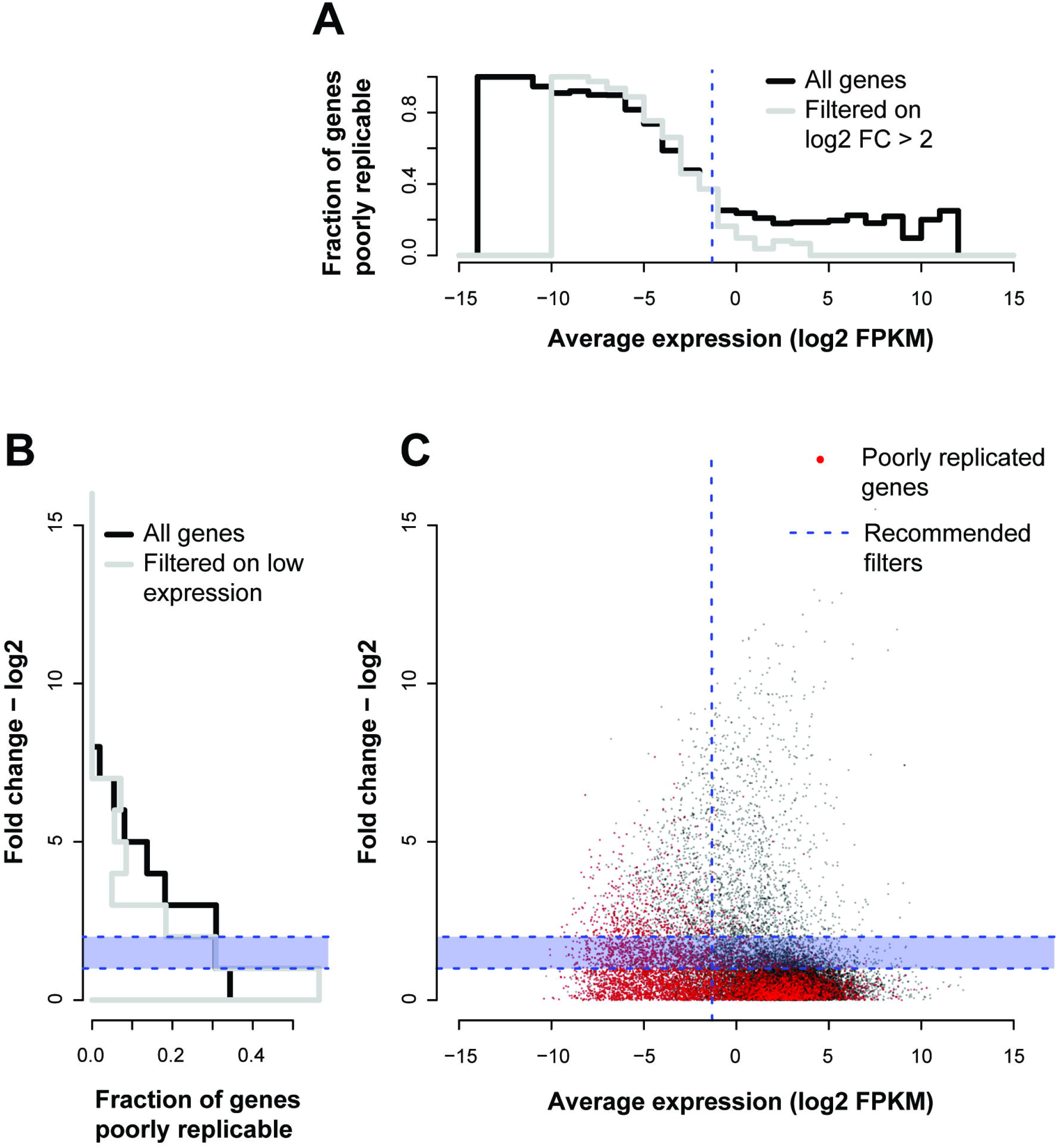
Low expression levels and high fold changes provide sensitive quality control. (A) Histogram of fraction of genes poorly replicable and filtered on mean expression (B) or fold change. (C) Plotting the average expression against the fold change to compare gene-gene replicability to the SEQC criteria. The red points are genes that were not well correlated (Pearson’s *r*< 0.9) with their replicates across conditions. The fraction of these red points across mean expression (log_2_ FPKM) is shown in the histogram in panel A, and the fraction of these red points across fold change (log_2_) is shown in the panel B. The recommended filters by the SEQC are shown by the dotted blue lines, across both mean and fold change. We see that the fraction of poorly replicated genes drops significantly at the recommended filters – i.e., discarding fold changes less than log_2_ 1~2, and discarding the lowly expressing genes (bottom 1/3^rd^). The grey lines show the histograms for a given measure (mean expression –A, fold change -B) contingent on the SEQC criterion for the other having already been applied.

### PART III: Using co-expression as a proxy for replicability

#### A co-expression heuristic to estimate gene expression replicability

Motivated by our view that a gene’s co-expression across conditions with itself in a replicate is a useful view of replicability, we wished to determine how this could generalize to comparisons between different genes that are very tightly co-expressed or even stoichiometrically co-regulated. In the same way that a gene should be well correlated with itself across conditions, we hypothesize that there are genepairs whose functional relationship is so specific, that the pair can be treated as pseudo-replicates of one another. These pairs will show similar expression profiles and where that does not occur, we can infer poor replicability within a given data set. We call these pairs of genes “housekeeping interactions”. Selection of these pairs is described in further detail in the **Materials and Methods** section “**Selecting co-expressed pairs**” while much of the remainder of this paper provides evidence that these pairs serve as an extraordinarily sensitive means to detect disruption within data.

Proteins in complexes have been found to be highly co-expressed, such as in the proteasome [25,26], and these genes respond to changes in the cell or tissue in a tightly regulated way. This stoichiometry – where parts of a protein complex are required in fixed quantities - can be used as a useful filter to obtain candidate genepairs. We thus derived this set of gene-pairs - our housekeeping interactions - by extracting the top co-expressing partners from a large scale meta-analysis of co-expression data [27], and those that were annotated as protein complexes. That co-expression data is separate from the remainder used in this work (and thus may be thought of as training data for the analyses conducted here). We note that while no individual pair is guaranteed to be correct, our principal interest is in enriching for tight co-expression so that aggregate signals are detectable.

The top co-expressed pairs in the housekeeping interaction list are between a unique set of 1,117 genes (i.e., some genes occur multiple times). We observe that these genes are enriched for cellular functions related to the cell cycle (e.g., GO:0051301 cell division p<4.82e-89) or a cellular structure (e.g., GO:0000776 kinetochore p<4.91e-40) amongst similar others (**S1 Table**). We expect enrichment for these functions primarily because we have selected for genes that intersect with those annotated as protein complexes. We also suggest that these basic biological functions are plausible as characteristic of the set of genes with housekeeping interactions which will remain under regulatory control, even once subject to experimental perturbation (i.e., some subset of the pairwise relationships will be held constant or necessarily co-disrupted). We also consider pairs defined by strongly conserved co-expression across species as a secondary test set and a set defined purely by protein interaction as a third set.

#### Modelling the impact of perturbations on co-expression to estimate gene expression replicability

For any of these tightly regulated gene-pairs, we expect that variation in the expression of one gene should match the variation in expression of its co-regulated partner, and it is this matching which we evaluate in the following. Going through our methodology in steps, a gene’s expression levels in an experiment along with the same data for the gene’s co-expression partner gives two gene expression profiles (**Fig. 6A**). It would be expected that if the expression data for these genes shows some differential signal across samples (i.e., biological differences), these two profiles would show a high correlation across the samples (**Fig. 6B**). However, if one or more samples have low precision in its estimate of the gene expression, this will weaken their correlation. Of course, for the genes to serve as equivalent to replicates, they must be subject to primarily independent sources of error. This is clearly not wholly true – minimally, they’re drawn from the same sample – but co-expression which is conserved, defines a protein complex or is seen in many independent experiments seems unlikely to be due to shared artifacts (although we evaluate this).

**Fig. 6.**
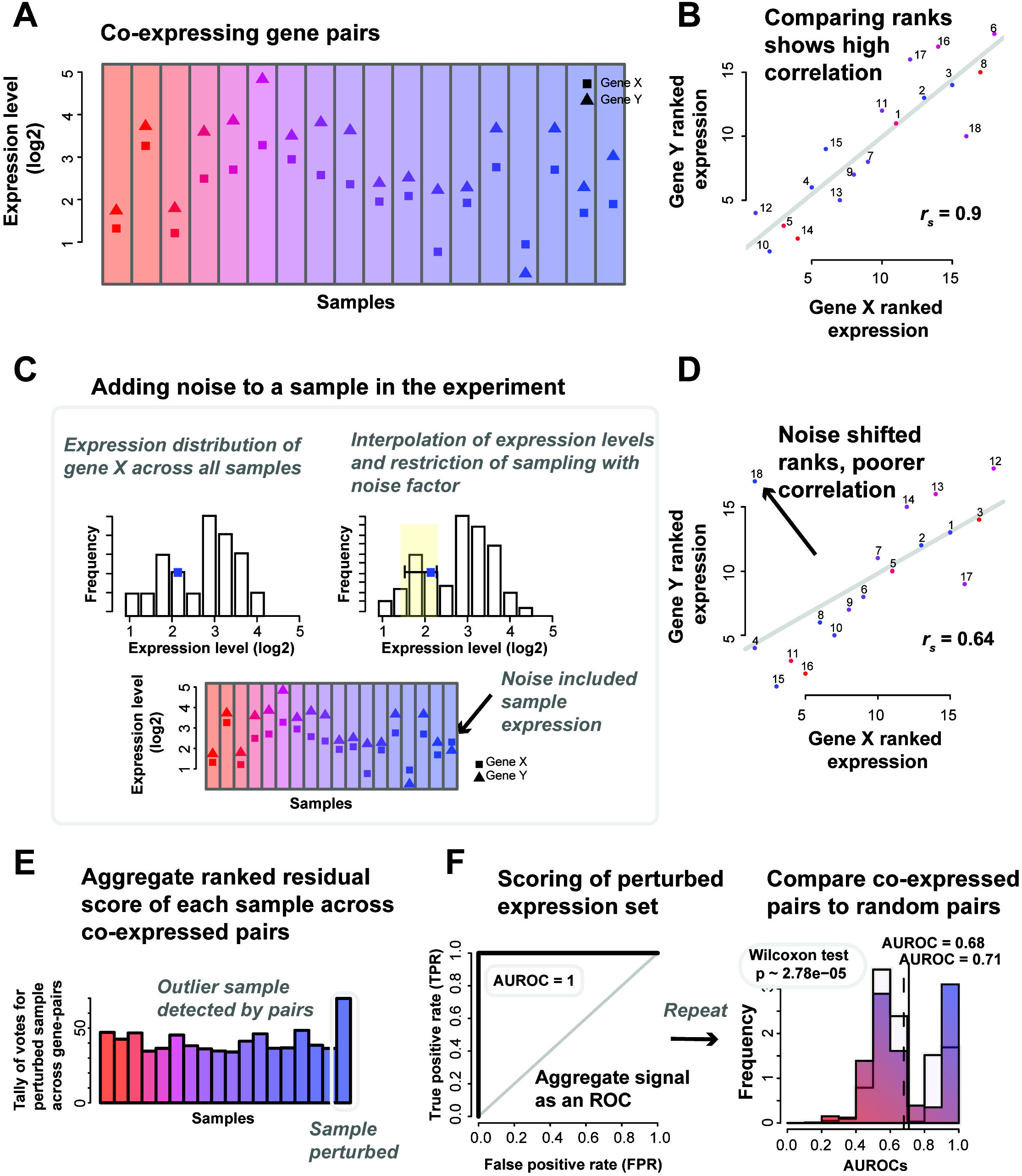
Co-expression perturbations to pick out samples with high variation. (A) Expression levels of genes X and Y. (B) Gene X and Y show good correlation (Spearman’s *r_s_* =0.9). (C) If we add noise to one sample, we see a shift. The noise model is described further in the Materials and Methods section. Briefly, for each gene in a sample, we select a new rank for it to take relative to its expression in other samples, thus sampling from within its empirical distribution. The new rank is limited to one close to the original value, as defined by the noise factor. (D) The noise added to the sample has caused it to be an outlier that is disrupting the co-expression indicated by the otherwise good linear fit. The residuals of the points scores the sample (regressing from the line of best fit), and the (E) scores in aggregate allows us to draw an (F) ROC and calculate an AUROC, testing how well we the outlier was detected. We also calculate a p-value (Wilcoxon test) to compare the distributions of the average AUROCs of the co-expressed pairs and an equal number of random pairs.

We quantify the strength of the co-expression relationship by introducing “noise” into the experiment through the perturbation of the expression levels of the gene in a particular sample (**Fig. 6C**). In our case, the noise added is a particularly subtle form of modification since we only nudge the gene’s expression by a slight amount to another value identically sampled from its expression distribution. Thus, in our case, 10% noise (perturbed) means that the original expression value is nudged up by enough to put the gene 10% higher in the rank that its new value takes relative to all of the expression values observed for that gene, the empirical distribution of its expression. Thus, if we rank standardize the data, the new value is sampled uniformly from samples ranking within 10% (out of all samples) of the original sample’s rank. Note that this is not a 10% change in its rank but a 10 % change out of the total. Linear interpolation between adjacent sample ranks allows us to sample continuously. Put simply, a gene’s expression level in one sample is replaced by its expression level in another sample. If that sample was the one with the closest expression level, the noise is low and if it is was the furthest, the noise is high.

Note that this is a subtle enough shift that it can never be detected from the gene’s expression alone; the distribution of expression values is essentially unchanged. If a small perturbation (i.e., low noise factor) is detectable, it indicates that the relationship between gene-pairs must be very clear (outliers are easy to identify, **Fig. 6D**). Equivalently, if a large perturbation is not detectable, it indicates an experiment with noisy data in which there are other samples that are still more aberrant than one where noise has been introduced.

Using the distribution of samples detected as outliers for each gene-pair (**Fig. 6E**, see **Materials and Methods** section **“Perturbation: the noise model”**), we can predict the perturbed sample, and from this determine the area under the receiver operating characteristic curve (AUROC, **Fig. 6F**) to characterize how well all the gene-pairs (in aggregate) picked out the perturbed sample (see **Materials and Methods** section **“Perturbation: the noise model”**). Repeating this through all samples gives an average AUROC at a particular noise factor – where an AUROC of 0.5 means that the noisy sample was undetectable, while an AUROC of 1 means that the perturbed sample is always being correctly ranked as the poorest. Thus having a high ability (i.e., a high AUROC) to detect even low noise is indicative of good performance – and good replicability as we now define it- while experiments with weaker performance will exhibit little change in response to noise and have lower AUROCs. Systematically perturbing the samples by the addition of “noise”, and measuring the effect (the AUROC) each perturbation has on the gene-pairs in aggregate can be used to quantify the quality of the experiment. As an analogy, this is like yelling in a room; if the noise is undetectable, the data is noisy to begin with, whereas if even a small noise is easily detected, the data is ‘quiet’ (shows pristine co-expression). It’s important to note that this model provides a relative ranking of all samples for their degree of noise; it is not the equivalent of a model which attempts to detect ‘differential’ samples. Thus, AUROC performance ranking the perturbed sample does not map to a categorical judgment that a particular sample is detected as different; just that it is correctly ranked as the most perturbed sample.

#### Application of co-expression for noise estimation in reference datasets

To test our concept, we selected the BrainSpan reference set as it contained a large number of samples. Using the set of co-expressed gene-pairs we derived, we varied ‘noise’ levels and sample sizes and recorded the resulting AUROCs. First, we note that randomly sampling a new expression value in a given sample for each gene (from within that gene’s expression distribution) completely disrupted the co-expression at that sample within the experiment and we could perfectly identify the disrupted sample (**Fig. 7A**, 100% noise factor, i.e. random). And with no noise, the AUROC was 0.5, since no signal from the gene-pairs could be used to distinguish the ‘disrupted’ sample from any others in that case. In this reference data with 500 samples, 5% noise was enough to increase the AUROC to 0.73.

**Fig. 7.**
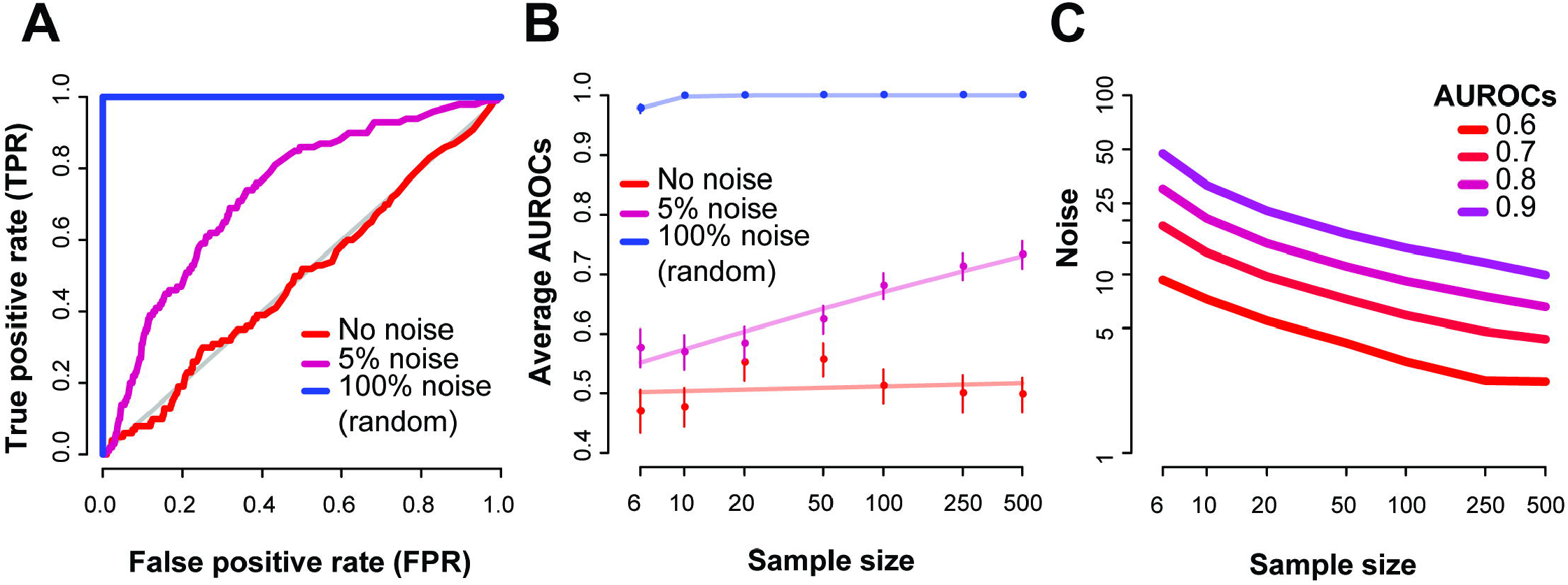
Replicates via co-expression pairs to extract the noisy samples. As an example, we use the BrainSpan RNA-seq dataset on 500 samples. We randomly added noise to one sample at a time at a given noise factor, and repeated this 100 times. (A) For each run, we generated an ROC, and show the average ROCs of these 100 runs, with the AUROC ranging between 0.5 and 1. Noise factor of around 5% was enough to disturb the experiments replicate, giving an AUROC of 0.73. (B) Average AUROCs varying across sample sizes and noise factors, with the standard error bar. Each point is an average of 100 repeats. The lines represent a linear fit of the data points. (C) Predicted noise factors for varying AUROCs shows the dependence on sample size and level of performance for the BrainSpan dataset.

We then systematically queried the space of different sample sizes versus noise (**Fig. 7B**), and observe similar trends. Low noise (<1%) across all sample sizes had average AUROCs close to 0.5, indicating that variation below that is undetectable in the dataset. While specific noise levels are shown in **Fig. 7B**, we could alternatively ask what level of noise is detectable to a particular accuracy (**Fig. 7C**). If we set the AUROC to 0.6 (i.e., low but significant detectability, equivalent to a Mann-Whitney p~1e-15), approximately 3.2% noise added to a sample (out of 100 samples) disrupts co-expression enough to correctly identify the sample at that level. This drops to 2.5% noise when the data has 500 samples, and increases to 4.1% for 50 samples.

To observe the general range of values in other experiments, we ran the analyses on a further 83 RNA-seq experiments, of varying sample sizes. The noise factors that would give an AUROC of 0.8 ranged between 3% and 40% and were inversely correlated to sample size (Spearman’s *r_s_*=-0.48, **S2A Fig**). This is not a surprising finding, however, it reaffirms that one of the most important influences of the quality and outcome of an experiment is the number of samples. Even so, for some smaller sampled experiments (<10), we recorded smaller detectable noise factors, similar to some of the larger experiments.

#### Robustness analysis of AuPairWise

Having demonstrated that the method detects subtle disruption, we wished to establish proper use and controls. First, we assess the relative efficacy of this method to what real replicates would measure (**Fig. 8A**). Instead of the subset of co-expressed pairs, we looked at all genes in the samples, paired to their replicate. As expected, replicates perform better than the co-expressed pairs (shaded lines versus solid in **Fig. 8A**) with true replicates allowing us to detect the addition of noise at a level ~1.25 times lower than co-expression replicates (averaged across all ROCs; e.g., if 5% noise is detectable at a particular AUROC with co-expression, a true replicate could detect 4% noise).

**Fig. 8.**
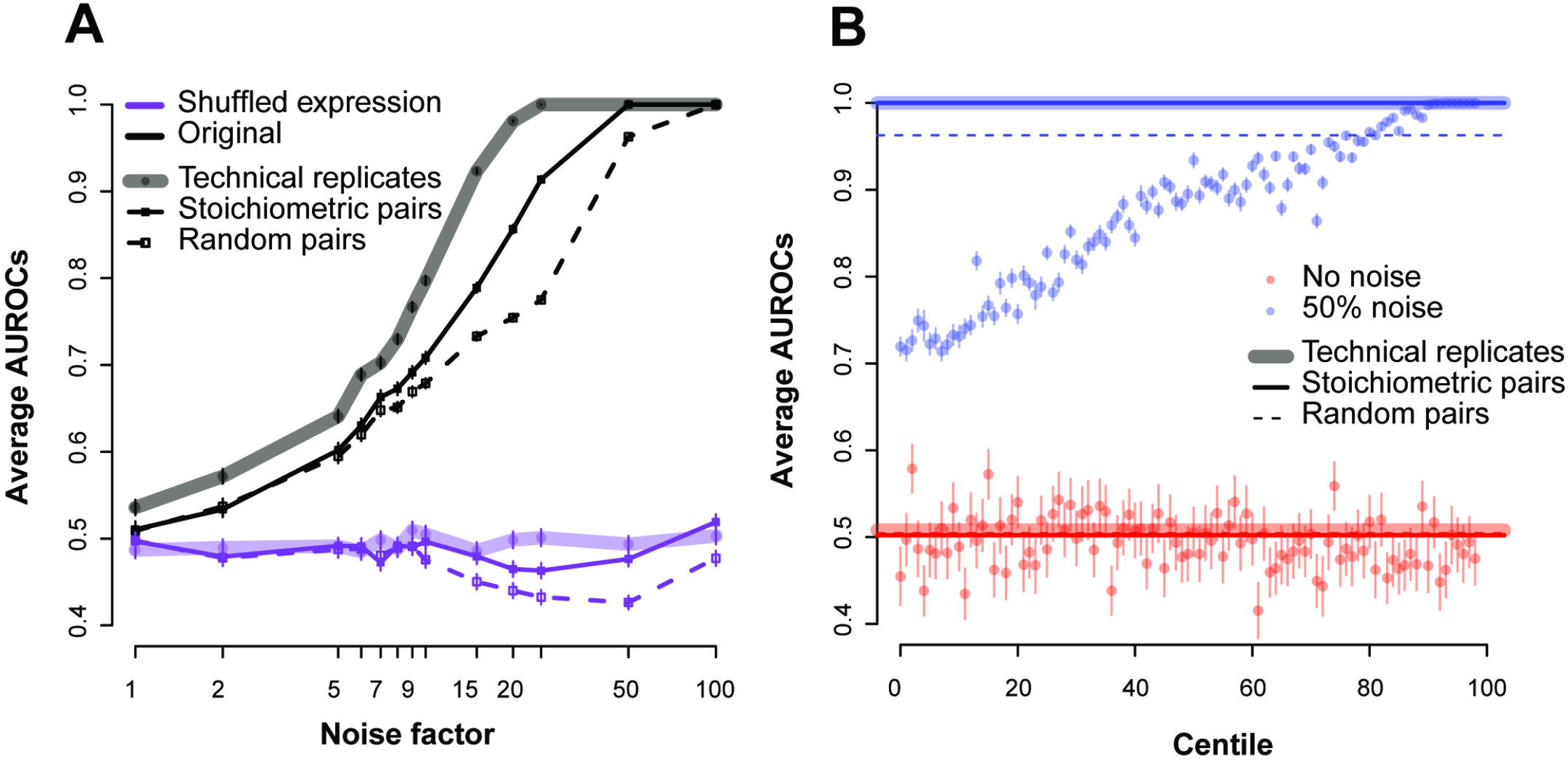
Estimating replicability using different gene-pairs. (A) Comparing the use of co-expressing pairs (solid), actual replicates (shaded) and random pairs (dashed) for the ENCODE dataset. Each point gives the average AUROC for detecting a noisy sample in the data across the varying noise factors. The sample size here was held constant at 18. The black lines are the original expression values, and the purple lines are those where we randomized the expression values (shuffling within an experiment). (B) Performance of the centiles for the experiment with and without the addition of noise. We only show the 50% noise factor. The replicates and stoichiometric pairs show very similar performance at this noise level (AUROC~1).

To better characterize these properties, we measured the performance of all possible gene-pairs (ranked from highly co-expressed to not at all). Dividing all the gene-gene correlations into centile bins, we measured the performance of these “centile pairs” under different noise factors (**Fig. 8B**). We expect that the stronger the correlations (higher centiles with better co-expression), the higher the average AUROC will be. We can see that it requires highly co-expressed pairs (in the 90^th^ centiles or greater in this experiment at 50% noise) to achieve performance close to the stoichiometric pairs, suggesting that the stoichiometric co-expression pairs are a particularly effective set to detect RNA-seq sample quality. Since these centiles are with reference to the maximum possible performance in this dataset, a perfect ranking with respect to them for our fixed set is not expected.

While we focused on high co-expression in protein complexes, we also wished to assess other plausible housekeeping interactions (**S3A Fig**). Our second set of gene-pairs consisted of highly co-expressed genes that were conserved across yeast and mouse (see **Materials and Methods** section **Selecting co-expressed pairs**). We also wished to assess a set of gene-pairs obtained independent of all expression data. We therefore looked at protein-protein interaction pairs that are part of protein complexes without regard to co-expression. Out of all the sets of pairs tested, the human stoichiometric co-expression pairs (i.e., those using PPI data and prior expression) performed the best, the yeast-mouse co-expressing pairs also showed good performance, while the purely protein-protein pairs showed moderate performance.

As a negative control, we selected sets of random pairs of similar number to the number of stoichiometric pairs, and once again ran the analyses. One striking feature of these results is that random pairs exhibit modest performance in all cases. Intuitively, one might think that random pairs could have some form of co-expression that would be detectable in aggregate, perhaps due to a few real and highly co-expressed pairs having been randomly selected or a more broadly distributed weak effect across all gene-pairs, not unlike the principle underlying correction for batch effects. Any factor other than biology which contributes to co-variation across samples could introduce this as an artifact and normalization confounds seemed to us a potential concern. Importantly, our method’s performance is not at all sensitive to changing the different normalization protocols (**S3B Fig**); however, a more finely calibrated assessment distinguishing technical artifacts of this type from biologically real variation would be to use the random gene-pairs as controls relative to our stoichiometric co-expression. The difference in performance between the two (**S4 Fig**), while not extreme, is actually quite stark: testing the performance distributions against each other yields a p-value of 2.78e-5 (Wilcoxon test, **S4B Fig**). Since technical variation should drive co-variation among random pairs and biology the preferential utility of stoichiometric pairs, we also report this statistic as a default within AuPairWise. Just as in differential expression it is important to have large differences for a subset of genes (biology) but not all (batch effect), so the same is true in our co-expression based analysis.

We repeated our analysis of the 83 independent RNA-seq experiments looking for purely technical co-expression as indicated by the performance of random pairs. In many of these experiments, even random pairs have good performance, and while this was generally very significantly less than the stoichiometric pairs (**S2B Fig)** only a very few experiments did not exhibit this potential technical confound.

We next assessed the robustness of our method to variation in the number of runs and selection of pairs (**S5 Fig**). For this analysis, we first varied the number of perturbations we ran for each noise factor analysis, and then calculated the variance of the AUROC. The noise factor has a strong effect on the variance, while increasing the number of runs, of course, lowers the standard error. We then varied the number of gene-pairs used in the analysis, and it appears that a minimum of approximately 500 or so pairs is enough to detect an aggregate signal.

One important caveat attached to the high performances we have reported is that our perturbation model has only perturbed one sample at a time. In reality, it is quite possible for the biological character of the data to be confounded with noise and, for instance, half the data to be of lower quality than the other half. For example, a differential expression experiment in which half of the samples were from a rare condition, hard to obtain, and for which worse quality control needed to be tolerated, would be one in which a lot of co-expression might be disrupted. Just as our single sample perturbation resembles leave-one-out cross-validation, we can conduct an identical experiment in 2-folds. We randomly sampled half the data from our ENCODE-based experiment and subjected each gene in each sample to an independent perturbation. Average AUROC’s for a given perturbation were 0.86 for 25% noise (significantly different than random pairs, Wilcoxon test p~6.65e-28, **S2 Table**). This comparably high performance also suggests that our modest class imbalance in the leave-one-out analysis is not seriously distorting assessment. Interestingly, one major difference in the 2-fold AUROC is that it does not go to 1 as the perturbation goes to 100%, instead reaching a value modestly below this (i.e., ~0.98, **S6 Fig**). Normally larger perturbations make it easier and easier to detect the altered sample(s), but when half of the samples have no co-expression due to the perturbation, it begins to impede the method. This is a fairly extreme state, however.

Finally, we considered whether our method could be naively applied to microarray data. Our recent work suggests that pseudo-replicates in co-expression data are dominated by artifacts in microarray data [27]. In particular, very high co-expression in microarray data is much more likely to reflect such artifacts (e.g., cross-hybridization) and occurs among low-expressing genes. While we caution that this analysis is only preliminary, applying AuPairWise to the equivalent microarray data set version of BrainSpan (across a common set of 495 samples, but performed on a microarray platform) also showed good performance (**S7 Fig**), yet the random pairs had almost equal performance (see p-values, **S3 Table**), implying a potentially greater role for normalization.

#### Guidelines and availability of AuPairWise

Our results indicate that using co-expression is a practical approach to predict the consistency of replicates across biological samples in expression experiments. To share this method, we have compiled our R scripts and make this available on github under the name AuPairWise, so named for its use of selected housekeeping interactions (github.com/sarbal/AuPairWise).

A summary schematic of the input and output is shown in **Fig. 9A**. The scripts are dependent on a few standard R packages and require expression data as input, with rows labelled with the set of genes detected in Entrez gene IDs format and columns labelled with the sample names. Given the expression dataset, a single run of the method outputs the average AUROC for a range of noise factors (which can be specified), along with the number of repeats. Further to this, an estimate quality control metric (“noise level”) is returned that is detectable over the threshold of 0.5 – i.e., this is the estimated amount of noise in the samples. As we wish to qualify the experiment based on biological properties, we compare the gene-pair results to those a similar number of gene-pairs chosen at random would do and return the p-value (Wilcoxon rank sum test, two sided) for each noise factor. When interpreting results from AuPairWise, we recommend reporting both the average AUROCs of the stoichiometric pairs, the random pairs, and the corresponding p-value (**Fig. 9B**). We suggest looking at a noise factor between 5%-25% (depending on sample size), with smaller sized experiments warranting higher noise factors for meaningful perturbations. For a given noise factor, a high performance on the stoichiometric pairs and low performance on the random pairs and therefore a low p-value is a “good” experiment. Low performance of stoichiometric pairs implies a poor run, and high performance on the random pairs implies there is systematic artifact in the experiment. We also recommend a minimal number of 10 samples for any of the analyses to be meaningful.

**Fig. 9.**
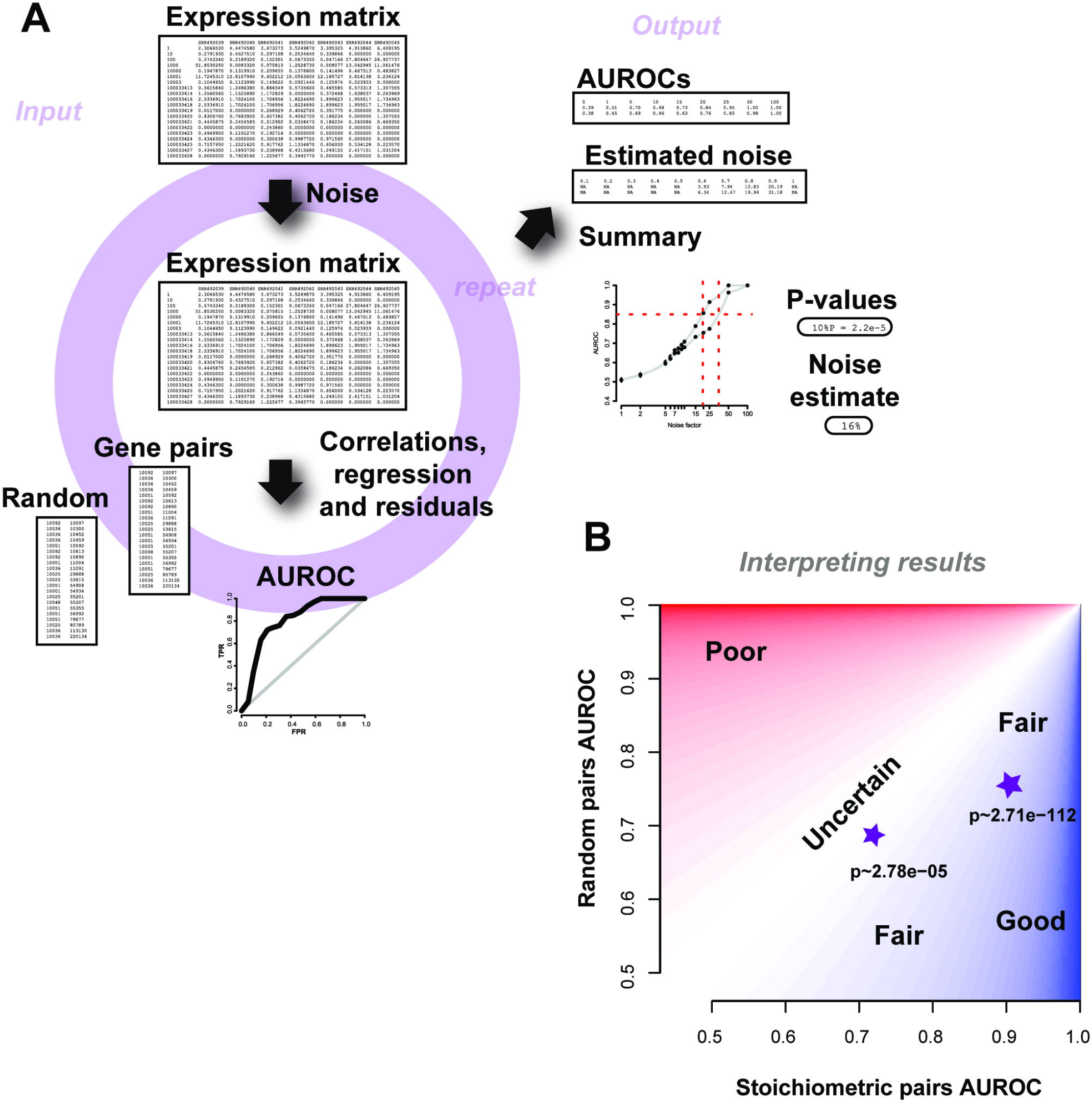
Schematic of the AuPairWise method and guidelines for interpreting results. **(A)** Input into the script is an expression matrix. Noise is added to a sample at random, and an AUROC is calculated based on how well the perturbation is detected by the gene-pairs. This is repeated for multiple noise factors, which then allows us to estimate the amount of noise required to significantly disrupt the experiment, which is used as our metric for replicability. The outputs are summary files with the AUROCs and noise estimates, along with the summary plot. (B) Guidelines for interpreting results. We plot performance of the stoichiometric pair against the random pair. Two toy examples are shown as stars, with their corresponding p-values beneath them. High performance (AUROCs) on the stoichiometric pairs and low performance on the random pairs for a given noise factor implies a fair to good experiment (dark blue shading). Any results that fall closer to the identity line are less certain and probably contain systematic noise (lighter regions), and are likely poorer experiments (red regions).

The current housekeeping interaction list is not perfect, but we argue that the list should retain, on average and in aggregate, a strong fundamental biological signal. Of course, if the experiment in question perturbs these pathways and protein complexes, it will be a useful set of gene-pairs to investigate. To this end, gene-pairs selected can be customized to exclude those subject to experimental perturbation, or those not measured. We have currently only made the human (*Homo sapiens*) datasets and pairs available, yet, as the pairs list can be modified, other species can be used, subject to further research to establish proper protocols and interpretation.

The current runtime for a single analysis (i.e., for 20 samples, 20,000 genes, 2500 gene-pairs, across 10 noise factors and 100 repeats) is approximately ~5 hours on our cluster of 16 cores with processors running at 2.4GHz and this scales linearly (**S5 Table**). However, we recommend running 1000 repeats. Although not formally implemented in the method, runs can be parallelized and the performance AUROCs combined across these runs.

## Discussion and Conclusions

Recent analyses of RNA-seq have added a number of caveats to its use, as is to be expected with any new technology as it begins to become mature [6,9]. Because of this growing appreciation of limitations in RNA-seq, we think a reconsideration of some of the early enthusiasm is timely. In particular, one of the most important technical claims made about RNA-seq – that it was highly replicable – was based on reproducing approximate transcript abundances. Instead, we have argued that we mostly should care about condition-dependent *changes* in transcript abundance. This is much harder to measure and much likelier to depend on experimental conditions. Consequently, it’s valuable to have a method which can measure the expected replicability of a performed experiment and it is this gap our analysis tool, AuPairWise, seeks to fill.

Quality control in RNA-seq is still a major concern, in particular when final biological outcomes are not replicated. At the heart of any quality control scheme is a ‘check’ for some expected class of result. For example, one might expect most genes not to be differentially expressed [28] and correct data to ensure that this is true. Our method hinges on an expectation that a particular subset of genes will show correlated expression. This is among the most well-validated observations in the analysis of expression data [25] and also one which has comparatively little impact on differential expression results. We have, nonetheless, taken unusual pains to ensure our method will not lead to overfitting, including assessing a variety of co-expression signatures, allowing customization of the pairs chosen for assessment, and not implementing our method as a correction.

Approaches related to our own are touched on in recent literature, even though most quality control in RNA-seq is focused on batch corrections of some sort. For example, in their analysis of single-cell data, Buettner et al. [29] exploit co-expression among cell-cycle genes to perform a batch correction based on cell-cycle as a latent variable. While this is most readily understood as a variation on conventional batch correction, it resembles our own technique in exploiting function in the form of known co-expression to determine a covariate associated with noise. Indeed, had that analysis generalized across many functional classes and then modelled the predictability of noise as a means of quality control (rather than as a correction), it would resemble our own method. Another related approach is the L1000 analysis platform which measures gene expression over only a small subset of genes and then uses learned relationships based on co-expression to infer the rest [30]. While this is exploiting an effect similar to that in our analysis, we rely on such relationships to infer a single measure of quality control; we are never too reliant on any one being accurate. To literally infer unknown expression from such relationships seems to us likely to lead to overfitting. A less extreme approach similar to the L1000 system is used in imputation of expression, where the expression of genes missing from a sample in the data are estimated by using the expression of genes that are co-expressed [31]. There, the expression profiles of the co-expressed genes provide the expression of the missing sample. In our case, prior knowledge as to which gene should be co-expressed is used not to correct the data, but to determine if that sample has correct values. Finally, because co-expression is used to infer function in networks, its utility for doing so in any given data can be used as a form of quality control (e.g., [32]), but this lacks any quantitative mapping to differential expression.

The perturbation model we have used, while simple and fairly close to conventional ideas such as bootstrapping (plus constraint and interpolation), is not common within RNA-seq analysis. We suggest that even outside of our specific application, a strongly data driven sampling distribution for gene expression is likely more desirable than more theoretical approaches based on modeled expression values. The observation of subtly erroneous normalization has been previously discussed as a problem for co-expression analyses [33]; this result is similar to our own that random pairs show some performance. In particular, we do consider it critical to report both random and real pairs for performance since it is the relative performance of ‘real’ pairs that indicates the experiment is conforming to expectation; just as in differential expression, pervasive effects across the genome are not evidence of high biological signal, but rather high technical signal. It is worth noting that co-expression network analyses are mostly robust to these normalization effects since they usually depend only on the relative rankings of correlations between all gene pairs and not on the significance of the values directly.

Our method is not without limitations. As it requires there to be a measurable amount of variance within the experiment, it will be difficult to calculate co-expression in an experiment that has no conditions for expression to vary across. The number of samples that make for meaningful co-expression will also need to be followed [27], although this seems more feature than bug to us, since it is, in fact, true that more data permits for more replicable results. We also suggest that the co-expressed pairs themselves are subject to review and reanalysis. As mentioned, we have tried our best to not overfit to current data, but it does rely on the available information present in co-expression data and gene annotations. As more data does become available, and our knowledge base on stoichiometric pairs and the biological implications behind these expands, the housekeeping interaction pairs will need to be refined. Of course, even regular housekeeping genes lists are by no means without argument despite decades of study. We also have ignored the transcript specific level of replication; this admits to clear expansion using transcript specific pairs. One final caveat is that because this method is not a correction of any type, the results do not pinpoint particular genes that are misaligned, but rather indicate that the whole experiment does not conform to expectation.

While previous work hints at the value of co-expression for quality control, we think it has been hampered by a poor sense of what precisely quality control should be controlling. We think our analysis and particularly our recapitulation of SEQC’s observational heuristics from more basic principles help guide quality control by attaching a more precise meaning to ‘replication’. While RNA-seq’s signal-to-noise ratio may be less idealized than early analyses implied, our ability to assess where noise *does* occur in RNA-seq appears to be much higher. By providing a common standard for that assessment, we hope AuPairWise will permit experimentalists the opportunity to refine their experimental design and validate their success in doing so.

## Materials and Methods

### Datasets

To illustrate our analyses, we used data from publically accessible RNA-seq experiments where we define an experiment as a set of samples. The majority of the work and analyses were performed using the following datasets: ENCODE (GSE35584) [34], Marioni et al (GSE11045) [21], Xu et al (GSE26109) [22] and BrainSpan [35]. The ENCODE dataset and the Marioni et al. dataset were downloaded from SRA and converted to fastq files using “fastq-dump” from the SRA Toolkit [36]. Following quality control using the fastX tools, we mapped the reads using bowtie2 [37] and estimated the RPKM and FPKM expression levels using RSEQtools [38] and Cufflinks [39] respectively, and the GENCODE annotations file [17] (version 18). We used the processed data provided in the Xu et al. supplement and the online Brainspan data. For our assessment of publically available data, we looked at those accessible both via Gene Expression Omnibus (GEO) and Sequence Read Archive (SRA). We used the URL from the GEO browser [http://www.ncbi.nlm.nih.gov/geo/browse/?view=samples&display=500&type=10&tax=9606&sort=type], filtering on human samples and those in SRA as of January 2015. Of the resulting 39K samples, we then filtered on samples only in a single experiments (unique to one GSE ID, close to 1.4K experiments), and then calculated the sample size for each of these. For a final follow up, we then obtained 83 RNA-seq experiments from the Gemma database [40] that were processed using RSEM [41]. These experiments varied in experimental design, sample size, read depth, sample type and conditions.

### Measuring replicability: correlations and sample estimates

We define an experiment as an ordered set of expression values for genes across multiple samples, with each sample defined as belonging to some class of biology from which RNA can be extracted. A replicate experiment is defined as an experiment across samples with identical class memberships, but varying on one of three other levels. This could be varying at a technological level (same samples, but run on a microarray), technical level (same samples, but a different run of the same machine), or a biological level (same conditions, tissue, cell, organism etc). The ENCODE dataset were purely technical replicates. To measure the replicability of an experiment in sample-sample space, we took the expression levels of an experiment, and calculated the Spearman (*r_s_*) or Pearson (*r*) correlation coefficient for each sample to its replicate (see Equation 1). Thus, we get a single correlation value for a sample, and multiple for an experiment. In our second measure of replicability, we are assessing the degree to which changes in expression for the individual genes are replicable. For this, we take each gene’s expression value across samples in one experiment, and the replicate values (i.e., the technical replicates), and calculate the Spearman or Pearson correlation coefficient of these two expression profiles for the same gene. For this measure, we get a value for each gene, and a distribution of correlation values. So, for a gene *i*, the gene-level replicability (*corr_i_*) is equal to the correlation (Spearman’s *rs_i_* or Pearson’s *r_i_*) between the expression values of the gene in experiment *X* (i.e., *X_i_*) and the replicate experiment *Y* (i.e., *Y_i_*), across *n* samples.

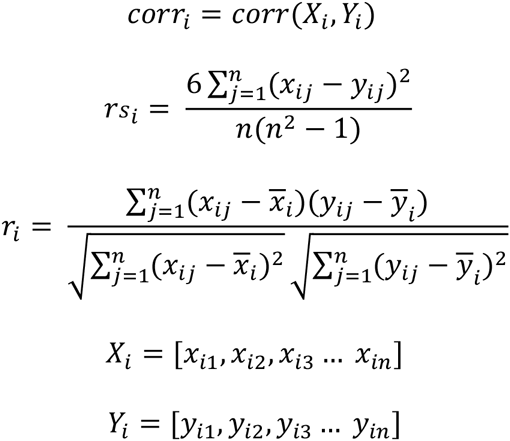

**Equation 1 Gene-gene correlations as a gene-level replication metric**

These gene-gene correlations each have an associated p-value that is calculated by either estimating a z-score (for a normal distribution, Equation 2) or a t-statistic (for a *T*-distribution, Equation 3), where *r* is the correlation value and *n* the number of samples in the experiment

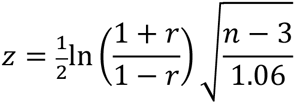

**Equation 2 *z*-score estimate**

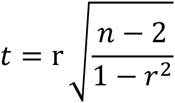

**Equation 3 *T*-statistic estimate**

One interesting terminological issue which overlaps with these two senses of replicability is the use of the term “RNA-seq experiment”. It is not uncommon within the literature to regard a single data sample as an experiment under the viewpoint that the different expression levels of different genes are then different measures of expression level. This would be close to the sample view of replicability in which even two samples would permit an estimation of replicability across those measures. In this work, we reserve the term “RNA-seq experiment” for data generated across multiple samples, typically to test some conditional variation. In such data, each gene will have multiple expression values and so gene-level replicability is meaningfully assessable.

We used the R functions **pnorm** and **pt** to estimate the p-values (for the z-score and t-statistic respectively). After adjusting for multiple tests with **p.adjust** in R, we then calculated the number of genes replicated as the number of genes with a significant correlation value (q<0.05). To estimate the number of samples required to get 90% of the genes replicated, we assumed that the range of correlation values would remain the same, but varied the number of samples *n* (within the t-statistic estimate equation), and recalculated the adjusted p-values. This gives us, for each *n*, the number of genes that will be significantly correlated, assuming the distribution of correlation coefficients is the same.

To compare our measures of replicability to the MAQC filters, we calculate the fold change (**Equation 4**) and the average expression of the ENCODE experiment. We took half the samples as untreated and the other half as treated, as per the experimental design (see GSE35584), and calculated the fold change as the absolute value of the ratio of the averaged expression levels between those conditions.

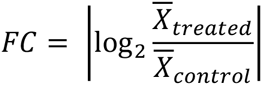

**Equation 4 Fold change calculation**

### AuPairWise: replication through co-expression

The overall idea behind the AuPairWise method is to leverage the biological signal of co-expressing pairs as a measure of replicability of an experiment. Therefore, the method requires a set of co-expressed pairs (section **Selecting co-expressed pairs**) that can be used as the biological indicator, and a noise model (**Perturbation: the noise model**) to determine the strength of these pairs. As a general overview, the nature of the test is a way to calibrate the noise within the system, as it attempts to detect outliers where we have purposely generated them. The more detectable noise is, the lower the original noise in the data (without the perturbation). As there are outliers in the real data, those samples are precisely the ones where we can add a substantial amount of noise without detection. Thus, when we perturb out system with noise, we are making a statement about the data prior to the noise being included. The degree to which perturbations can go undetected is the degree to which the experiment is noisy.

### Selecting co-expressed pairs

We generated a list of co-expressed pairs of genes that are required, in a similar way to housekeeping genes, to be expressed at stoichiometric ratios. First, we took the top 1% of genes that are reciprocally co-expressed in an aggregate microarray co-expression network. We similarly took the top 1% co-expressed genes from an aggregate RNA-seq co-expression network. We then took the intersect of these two gene-pair sets. To limit our set to what we believe are non-condition specific pairs, we took all the gene-pairs that were annotated as protein complexes in the Gene Ontology (GO [42], GO:0043234). With this, we were left with 2,669 genepairs, between a total of 1,117 genes. We also generated a similar list of co-expressed pairs conserved in yeast and mouse. We took the top 1% co-expressed pairs from an aggregate yeast co-expression network that we had created from 30 experiments across 966 samples on the Affymetrix Yeast Genome S98 Array (GPL90) platform. We also then took the top 1% co-expressed pairs from an aggregate mouse co-expression network (30 experiments across 1,575 samples on the Affymetrix Mouse Genome 430 2.0 Array, GPL1261). We then took the intersect of these two lists, whereby we had homologs for both genes in the pair and were left with 352 genepairs. Another gene-pair list we used was the protein-protein interaction set from BIOGRID (version BIOGRID-ALL-3.2.106, June 2014). Once again, we filtered this list down to 7,731 gene-pairs (interactions) by specifying that the interaction was a physical interaction, from Affinity capture – mass spectrometry experimental systems, and only genes associated with the GO protein complex term (GO:0043234).

In order to better characterize what GO terms were enriched in our human co-expressed gene-pairs set, we ran a simple gene set enrichment analysis using the hypergeometric test (**phyper**), correcting for multiple hypothesis testing using the Benjamini-Hochberg adjustment (**p.adjust**) in R [43].

### Perturbation: the noise model

We define an experiment as the expression levels of all genes across all samples for a range of conditions. Samples are not constrained to biological replicates, but should contain at least two conditions. Using the expression levels of each gene-pair, we calculated a score based on the residuals of the linear regression model fitting the points. Each point is a representative of a sample and its “expression replicate” (co-expression partner). Because we expect the expression levels of the genes to be “replicated”, there should be high correlation (the gene correlations across samples we defined earlier), and thus the residuals should be small, and “outliers” few and random. We rank these residuals, thus giving each sample a score. If a sample is perturbed, it should on average score the worst across all genes. We illustrate this in **Fig. 6**, and in more detail in supplementary **S8 Fig**.

In order to measure how robust the experiment was to noise, we then perturbed the system by randomly selecting a sample and added a relative amount of noise to all the genes in that chosen sample. We used a perturbation model that was a function of the dynamic range of a gene and the sample size of the experiment, such that a noise factor of 10 meant for an experiment with 100 samples the gene would vary within 10% of that gene’s expression values (as measured by the current experiment). First, we rank the expression profile of that gene (i.e., the gene’s dynamic range) across the samples. We then select a sample to perturb at random (**S8A Fig**). As we are perturbing each gene’s expression level for that sample based on its dynamic range, we need to ensure that the expression value is within the measured range of both the experiment and the samples. To do this, we order the gene’s expression values and interpolate the values between them to generate our sampling distribution (**S8B Fig**). We then calculate the rank shift for the given noise factor (i.e., for a 100 samples at a noise factor of 10%, we select a rank shift between 0-10), sampled from a uniform distribution (**S8C Fig**). At this new rank, we estimate the new expression value for that gene based on its dynamic range (**S8D Fig**). We then remap the expression value to be within the samples expression range (standardization). We used this model to ensure that the expression values remained within the empirical distribution of the gene and the expression distribution of the sample. Our expression experiment now has a perturbed sample (**S8E Fig**) with expression values within its original measurements. To be consistent, we defined random as a noise factor of 100%, and no perturbation as 0%. Noise is added in this way to all genes in that one sample, independently. Once all the genes are perturbed in this way, we then calculate the effect of this perturbation on the co-expressing pairs. We rank the expression profiles of each gene across the samples, and calculate the studentized residuals of the linear regression between the two genes. The score for each sample is simply the ranked residuals, whereby samples which deviate the most from the line of best fit rank the worst. With the aggregate scores from the individual linear regression model for all the genes pairs, we generated an ROC curve and calculated the area under the curve (AUROC). We also note that the AUROC is equivalent to the U statistic of the Mann-Whitney test (**Equation 5**), and can be calculated as such, where *P* is the rank of the held-out positive, *TP* the number of true positives, and *TN* the number of true negatives. In this case, we have *P* equivalent to the rank of the perturbed sample, *TP* to be 1, and *TN* to be *N*-1, where *N* is the total number of samples. If ranked correctly (i.e., *P = TP+TN = N*), we get an AUROC of 1. If ranked randomly, the AUROC value ranges between 0 and 1 and on average will be 0.5.

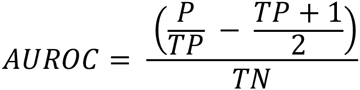

**Equation 5 AUROC calculation**

We used the AUROC as a measure of the performance of the experiment.

## Validation of pseudo-replication

As a proof of principle method for experiments where there were replicates available in the experimental design, we used the replicates and looked at every gene-paired with itself. In this case, we took the residuals of the gene replicates from the linear model, aggregating across all the genes (paired with their own replicate), and generating a ROC and calculating the AUROC. As described previously, we varied the amount of noise, repeating and permuting across the samples.

For the two-fold validation, where we perturbed more than one sample, we performed the same experiment, selecting at random half the samples and shuffling the expression values of the genes according to the noise factor. In this case, we shuffled each sample independently, so that the shuffled samples did not affect the values of the remaining samples to be shuffled. The AUROC was calculated as above, and the experiment was repeated across the different noise factors, and across different random set of samples.

### R scripts on github

For further information on how to the use the AuPairWise method and obtain the metrics, see the github repository (github.com/sarbal/AuPairWise). We have briefly summarized the code below (**Equation 6**).

~~~
For a given noise factor
   Chose a sample
      For each gene in this sample
            Add noise to gene’s expression
   Remap sample to original expression levels
   For each gene-pair
      Calculate line of best fit
      Calculate residuals for each sample pair
      Rank residuals to score each sample
   Aggregate ranks scores for each sample
   Calculate AUROC using aggregate scores
   Repeat
   Return average AUROC
~~~

**Equation 6 Pseudocode for the AuPairWise method**

## Acknowledgments

We would like to thank Sanja Rogic for assistance using the Gemma RNA-seq data and Paul Pavlidis for helpful comments on the manuscript.

## Supplementary Information

**S1 Text Additional text, data location, script description and references**

**S1 Fig Density of mean gene-gene correlations between replicates before and after filtering**. (A) For the fold change values, we see that for the recommended FC cutoffs (the blue dotted lines), the mean correlations are low and increase the further we move (black line). Removing low expressing genes shows even higher mean correlations (grey line). (B) For the average expression values, once again, we see that for the recommended cut-off, we have higher mean correlations (black line), that are improved even more with the inclusion of the fold change filter (grey line).

**S2 Fig AuPairWise noise metrics on RNA-seq experiments**. Here we took 83 RNA-seq experiments and ran the AuPairWise method for 100 repeats across a range of noise factors. (A) The density plot (solid line) shows that most experiments fall within the 19% noise factor range as assessed to cause an AUROC shift to 0.8. There is a dependence on sample size of course, with larger experiments requiring smaller noise perturbations (data points in figure). (B) Significance of difference in performance AUROCs for biologically related genepairs compared to random pairs for all the experiments. The geometric mean of these p-values is 4.67e-06. For comparison, the ENCODE reference dataset with 18 samples sits with a p-value of 6.52-e14.

**S3 Fig Average AUROC using alternative gene-pairs and different expression normalization methods**. (A) The co-expressed pairs (shown in red) outperform the protein-protein pairs (brown), the conserved yeast-mouse co-expressed pairs (pink) and random pairs (blue). These are the results of 1000 repeats. (B) Using different normalization methods on the expression data has a small effect on the performance AUROCs of the stoichiometric pairs. Here we compared the effects of RPKM (brown), TPM (blue), TMM (orange), VST (red), and simply ranking the expression data (green). These are the results of 1000 repeats.

**S4 Fig Distribution of the gene-pair AUROCs for different noise factor parameters** The distributions are for 1000 runs of the method on the same dataset, varying the sample being perturbed. The colored distribution is the co-expressed pair performances, while the lighter distribution is the performance of an equal number of random gene-pairs. The solid line is the average AUROC for the co-expressed pairs, the dashed for the random pairs. The p-value is the significance of the difference between these two distributions (Wilcoxon test, two-sided). Panels are for different noise factors (A) No noise added (p~0.85), (B) 10% noise (p~2.78e-5), (C) 25% (p~2.71e-112) and (D) 50% (p~2.76e-153). We see that although the average AUROCs may be similar, the distributions are significantly different.

**S5 Fig Variance versus parameter choices in the AuPairWise method**. Here we show the standard error (SE) of the performance AUROCs of the stoichiometric pairs as a function of the parameter choices on the ENCODE dataset, holding the sample size constant. (A) First we calculate the effect of the number of runs on the analysis, by varying the number of repeats from 10 to 1000. The SE decreases as the number of runs increases, as expected, with the noise factor having a small effect on the SE. (B) Varying gene-pair numbers (between 10 and 2500) shows a consistent SE only when 500 or more gene-pairs are used. To note, all variance is below 0.01 for almost all these parameter choices, indicating the results are quite robust. (C) We show the performances (average AUROCS) as a function of to the average number of genes within the sampled gene-pairs (and mapping between average number of genes and gene-pairs in inset).

**S6 Fig Performance AUROCs for AuPairWise on ENCODE data with the 2-fold cross validation**. Using the ENCODE dataset once again and running the method 1000 times, we varied the number of samples perturbed in the experiment. The n-fold cross-validation (one perturbed sample, in red) outperform 2-fold cross-validation (half the samples perturbed, purple). The 2-fold validation does not have an AUROC performance of 1 at 100% (~0.98), likely due to the fact that a great deal of co-expression is lost, hence preventing the method from detecting any structured co-variation.

**S7 Fig Performance AUROCs for AuPairWise on microarray and RNA-seq versions of the BrainSpan dataset**. The co-expressed pairs (darker lines) outperform the random pairs (lighter lines), for both the RNA-seq (green) and Microarray (blue) datasets. But the microarray dataset has more similar performances between the co-expressed pairs and the random pairs, compared to the RNA-seq version. The microarray dataset also performs worse/close to the RNA-seq random pairs.

**S8 Fig Expansion of Fig 6C explaining the noise model** (A) Expression of gene X from experiment. (B) The first step is to order the expression levels, and interpolate between the samples to get a continuous distribution of expression. (C) Then with this distribution, we define the boundaries for the noise included expression levels for that gene. For instance, with 10% noise, the new rank is limited to the yellow rectangle bounds, which corresponds to an expression change, constrained between expression values of ~1 to ~2 (the orange rectangle). (D) We then pick the new expression level and standardize it. (E) This is repeated until all the genes in the selected sample have “noise” added to them.

S1 Table GO enrichment of genes in the housekeeping interaction gene-pairs

S2 Table Significance of difference in performance AUROCs for gene-pairs compared to random pairs of the ENCODE dataset.

S3 Table Significance of difference in performance AUROCs for gene-pairs compared to random pairs for microarray and RNA-seq versions of the BrainSpan dataset.

S4 Table Performance and significance of difference in performance AUROCs for gene-pairs compared to random pairs for 2-fold cross validation of the ENCODE dataset.

S5 Table Average running time of AuPairWise across different sample sizes and repeats for a noise factor

